# Multivariate and Regional Age-Related Change in Basal Ganglia Iron in Neonates

**DOI:** 10.1101/2023.07.05.547821

**Authors:** L. Cabral, F.J. Calabro, W. Foran, A.C. Parr, A. Ojha, J. Rasmussen, R. Ceschin, A. Panigrahy, B. Luna

## Abstract

In the perinatal period, reward and cognitive systems begin trajectories, influencing later psychiatric risk. The basal ganglia is important for reward and cognitive processing but early development has not been fully characterized. To assess age-related development, we used a measure of basal ganglia physiology, specifically brain tissue iron, obtained from nT2* signal in rsfMRI, associated with dopaminergic processing. We used data from the Developing Human Connectome Project (N=464) to assess how moving from the prenatal to the postnatal environment affects rsfMRI nT2*, modeling gestational and postnatal age separately for basal ganglia subregions in linear models. We did not find associations with tissue iron and gestational age [Range: 24.29-42.29] but found positive associations with postnatal age [Range:0-17.14] in the pallidum and putamen, but not the caudate. We tested if there was an interaction between preterm birth and postnatal age, finding early preterm infants (GA<35 weeks) had higher iron levels and changed less over time. To assess multivariate change, we used support vector regression to predict age from voxel-wise-nT2* maps. We could predict postnatal but not gestational age when maps were residualized for the other age term. This provides evidence subregions differentially change with postnatal experience and preterm birth may disrupt trajectories.

## 1.0 Introduction

The perinatal period is a time of unique and consequential brain maturation, during which significant events, like preterm birth and the onset of postnatal stimulation, can uniquely impact development, setting infants up for a cascade of brain-based changes that contribute to later trajectories. Both as a cause and consequence of brain development, this period is also marked by the development of increasingly sophisticated motor and cognitive behavior. The basal ganglia plays a significant role in both cognitive and motor development, receiving the majority of the dopaminergic innervations from cortex (Prensa et al. 2000; Antonopoulos et al. 2002; Voorn et al. 2004; Paraskevopoulou et al. 2019). While a growing body of literature demonstrates the importance of striatal development for these behaviors in later developmental periods (i.e., adolescence (Geier et al. 2010; Luciana et al. 2012; Larsen and Luna 2015; Shulman et al. 2016; Parr et al. 2021)), basal ganglia neurophysiological development in the perinatal period remains understudied, and the impacts of preterm birth and postnatal stimulation have not been fully characterized.

MRI has been used to non-invasively probe basal ganglia neurophysiology across development. Previous work has used MRI based measures of basal ganglia tissue iron, such as R2’, to probe physiology (Larsen and Luna 2015; Parr et al. 2022). Brain iron is highly concentrated in the basal ganglia, (Hallgren and Sourander 1958; Brass et al. 2006) and is involved in neural processes such as cellular respiration (Ward et al. 2014), mitochondrial energy synthesis, including the production of ATP, (Horowitz and Greenamyre 2010; Paul et al. 2017) and monoamine synthesis, including of dopamine (Youdim and Green 1978; Ortega et al. 2007; Youdim 2018). MRI-based indices of basal ganglia tissue iron have been shown to increase throughout the lifespan, (Adisetiyo et al. 2012; Larsen and Luna 2015; Hect et al. 2018; Peterson et al. 2019; Larsen, Olafsson, et al. 2020) and deviations in normative maturational trajectories have been linked to pathologies that impact the dopaminergic system, such as Parkinson’s disease (Zucca et al. 2017), Huntington’s disease (Ward et al. 2014), and ADHD (Adisetiyo et al. 2014). As an additional MRI-based measure, tissue iron can also be assessed indirectly using normalized T2*-weighted (nT2*) echo planar imaging (EPI) scans that are acquired during standard functional MRI (fMRI) protocols. These show similar age-related changes to MRI sequences designed to obtain direct indices of tissue iron (Peterson et al., 2019; Larsen and Luna, 2015; Parr et al., 2022). Quantitative Susceptibility Mapping (QSM) is an exemplar sequence and directly indexes iron. Briefly, field maps are collected to form a spatial map of magnetic susceptibility, which is then deconvolved to produce one voxelwise map reflecting the tissue properties, including susceptibility indicative of iron deposition (Liu et al. 2015). R2’ is another exemplar sequence, where R2 is equal to 1/T2, and R2’ has been shown to contribute to the development of frontostriatal circuitry in adolescence (Parr et al. 2021).

A positive association has been demonstrated between positron emission tomography (PET) indices of dihydrotetrabenazine (DTBZ), reflecting presynaptic vesicular dopamine storage, and MR-based indices of tissue iron (R2’) (Larsen, Olafsson, et al. 2020), suggesting that basal ganglia tissue iron reflects dopamine availability. Furthermore, MR-based indices reflecting tissue iron properties have been associated with changes in cognition throughout childhood (Hect et al. 2018) and adolescence (Larsen, Bourque, et al. 2020; Parr et al. 2022), including working memory and IQ (Carpenter et al. 2000; Darki et al. 2016; Hect et al. 2018). Iron concentration has also been linked to reward sensitivity and the development of frontostriatal connectivity throughout adolescence (Parr et al. 2021 Mar 2; Parr et al. 2022). Although we are beginning to understand how basal ganglia tissue iron contributes to adolescent brain development, less is known about the role of iron in cognitive function in early childhood development.

The majority of an infant’s somatic iron during the first four to six months of postnatal life comes from iron stored during pregnancy (Rao and Georgieff 2007). Breastmilk has a relatively low iron content in comparison to Food and Drug Administration (FDA) regulated formula, optimized for the needs of infants, and breastfed infants have lower serum iron levels in comparison to formula-fed infants (Dube et al. 2010; Thorisdottir et al. 2013; Clark et al. 2017). In infancy, decreased serum iron has been associated with poor performance on a range of cognitive tasks. Umbilical cord ferratin levels, a measure of iron accessibility for the fetus, have been shown to negatively impact cognitive and motor behavior (N=278), including gross motor skills, auditory comprehension, and attention, assessed when the children were 5 (Tamura et al. 2002). Interestingly, increased cord ferratin was also associated with decreased full scale intelligence scores, suggesting that iron overload or overall disruptions in iron homeostasis impact cognition (Tamura et al. 2002). In line with the idea decreased iron leads to decreased scores on early developmental tests, anemic infants in Costa Rica, some of which were fed a partial diet of cow’s milk, low in iron, scored lower on both the motor and cognitive portions of the Bayley Scales of Infant Development (Lozoff et al. 1987; Lozoff, Jimenez, et al. 2006). Lower scores on cognitive tests persist even after being treated with iron therapy, suggesting that iron is necessary for cognitive development during critical developmental windows, such as infancy (Lozoff, Beard, et al. 2006). Further, cognitive deficits including working memory, planning, and inhibitory control (Lukowski et al. 2010; Algarin et al. 2017) continue throughout adolescence, suggesting a role for iron early in life in setting long-term trajectories in cognitive function. Imaging in this group finds abnormal visually evoked potentials and auditory brainstem responses, as well as abnormalities in resting state networks (Algarín et al. 2003; Roncadin et al. 2007; Algarin et al. 2017). These results are substantiated by additional evidence from pregnancies with clinical factors indicating possible iron deficiency (e.g. maternal diabetes and hypertension), showing low cord ferritin levels at delivery are related to slower auditory brainstem responses in neonates tested 48 hours after birth (Amin et al. 2013). Further evidence related to iron and brain development comes from work demonstrating maternal somatic iron during pregnancy was related to infant brain volume (Wedderburn et al. 2022).

The perinatal environment may have significant importance for the development of infant tissue iron deposition. Despite the potential for disrupted development, limited work has characterized the deposition of basal ganglia iron during this period. One way to examine accrual with development is to contrast issue iron deposition with postmenstrual age, combining the length of gestation with postnatal age to obtain a measure of the total age of the infant. Ning et al. (2014) assessed iron accrual using postmenstrual age (37-91 weeks), and found age-associated increases, with iron increasing across the first year of life. However, postnatal age might also be studied in isolation to measure the unique contribution of the postnatal environment. Initial work has identified increases in tissue iron with postnatal age. Zhang et al. (2019)’s work incorporated a sample with N=12 infants. Using two sequences specifically intended to measure tissue iron, QSM and R2*, they found age-related growth between age bins, where the oldest bin was in early childhood. These results are replicated by work with R2* and QSM in 74 children, (6-154 postnatal months), finding age-related associations for the putamen, pallidum, and caudate in a sample that extends into childhood (Raab et al. 2022). Again, assessing only postnatal age, Ning et al. (2019) identified regional increases in basal ganglia iron within the caudate, pallidum, and putamen in 28 infants (<6 postnatal months). Interestingly, when they examined the age at which linear age associations could be identified, they found associations in the pallidum and putamen after 3 postnatal months, but age-related associations did not emerge until 6 postnatal months in the caudate. Together, these studies suggest that basal ganglia tissue iron development may be evident during the postnatal period, but further investigation is needed to examine change within the perinatal period, specifically change associated with the alteration of brain trajectories when moving from the prenatal to postnatal environment.

The work described above has either used postnatal age or postmenstrual age to model tissue iron deposition in the neonatal period. Characterizing age-related associations using these measures provides us with the structure to understand neonatal change, allowing us to examine change with the length of exposure to the postnatal environment or overall linear change with development. However, as described in Cabral et al. 2023, additional work is needed to assess how brain development is uniquely impacted by gestation and subsequently altered by going from prenatal to postnatal life, where the brain rapidly changes as a response to postnatal stimulation. The perinatal variability associated with this switch has not been fully characterized. To identify this variability, we need to assess both gestational and postnatal age as separate terms in linear models. This isolates critical periods of change that may steepen or level off in a way that is not visible when characterizing iron accrual as an equally ascending process across development. If exposure to maternal iron stores during gestation resulted in significant infant brain iron accrual, linear change would be significantly associated with gestational age. An association with postnatal age might be seen if postnatal exposures were primarily responsible for iron accrual. These shifts in iron accrual may be independent or the prenatal environment may exert postnatal influence, possibly through an infant’s somatic iron stores, acquired primarily in pregnancy. If the prenatal environment was influencing postnatal deposition, an interaction term in the model could be significant.

Our previous work early infancy (PMA at scan: range 39.71-48.57 weeks) examined these relationships, with separate terms for gestational and postnatal age (Cabral et al. 2023). We found that postnatal age was significantly associated with an indirect measure of iron in the putamen and pallidum, but not the caudate. Gestational age was not associated with our indirect measure of iron in any of the three basal ganglia subregions. However, our sample was largely of term equivalent age, with only one infant <35 weeks gestation, which could make it difficult to see an effect of GA. To adequately model the relationships between indices of iron and both gestational age and postnatal age, as well as their interactions, a wide range of gestational ages, including preterm infants, is needed.

Multivariate approaches can also be useful to elucidate change associated with critical portions of development. Examining change solely on a regional basis may mask subtle associations with gestational or postnatal age. Connectivity patterns strengthen across motor and cognitive systems in early development (Grewen et al. 2015; Gao et al. 2017; Li et al. 2019; Molloy and Saygin 2022), potentially at different rates (Fiske and Holmboe 2019; Cabral et al. 2022). The basal ganglia has significant connectivity with these networks. Differently maturing connectivity within the basal ganglia has the potential to influence subregional tissue iron deposition. Iron may localize where white matter develops (Connor and Menzies 1996) and dopaminergic processing is needed (Ortega et al. 2007). This need for iron may differ at the subregional level. For example, the head of the caudate may actively participate in the basal ganglia association loop (Grahn et al. 2008), while the tail has been found to have substantial connectivity with the visual corticostriatal loop (Seger 2008; Seger 2013). Visual processes may develop early, resulting in a potential increase in iron deposition on the tail of the caudate before the head. This voxel wise iron deposition in the neonatal period remains unstudied. Preterm birth has the potential to disrupt basal ganglia connectivity patterns (Lao et al. 2016), and motivate an independent analysis of gestational and postnatal age effects on voxel-wise iron accrual.

Here, to examine both regional and voxel-wise iron accrual in the perinatal period, we used data from the Developing Human Connectome Project, which includes both preterm and term infants (Gestational age range: 24.29-42.29), with a significant sample size (N=464). We modeled iron using a measure of indirect deposition derived from the T2* signal in the resting state fMRI scans (nT2*). Iron has a short transverse relaxation time (T2) because of its paramagnetic properties. Therefore, iron is inversely proportional to the T2* signal, with increased iron reflective of decreased T2*, which makes it suitable for quantification with rsfMRI. In univariate analyses, we modeled age-related change using independent terms for both gestational and postnatal age, as well as an interaction to assess the interplay between gestational and postnatal development. To assess multivariate change, we used support vector regression to predict age from iron deposition independently for gestational and postnatal age. This was done independently for the term and preterm infants.

## 2.0 Materials and Methods

### 2.1 Participants

Data was obtained from the second data release of Developing Human Connectome Project (dHCP), which contains data from 512 infants. Some of these infants were scanned twice, and, if this was the case, we included only their first scan in the analysis, resulting in data from 464 unique individuals. Participants had a wide range of gestational ages (GA) and postnatal ages at the time of scan. Here, we were interested in early preterm births, and we defined prematurity as gestations that were less than 35 weeks to isolate potentially severe cases. The rate of NICU admissions increases with the degree of prematurity, with some work suggesting NICU admission rises almost 50% from 35 to 34 weeks (Loftin et al. 2010), with other work (N=3229) suggesting increases from 5.2% at 35 weeks to 34.5% at 34 weeks (Habli et al. 2007). Similar patterns are observed for respiratory distress, a common symptom of prematurity (Loftin et al. 2010) ((Habli et al. 2007). Thus, to isolate the potentially severe consequences of prematurity, we used < 35 weeks as a cutoff. The final sample consisted of N=84 preterm (GA Range 24.29-34.86 weeks; Postnatal Age Range 0.286-17.14 weeks) and N=380 term (GA Range 35.14-42.29 weeks; Postnatal Age Range 0-8.71 weeks) infants included in the analysis.

### 2.2 Scan Acquisition

Infants were scanned during natural sleep using a 3T Philips Achieva scanner. Scans for the dHCP have been described in detail elsewhere(Fitzgibbon et al. 2020) and include both a T1 and T2 weighted image for segmentation and normalization to the anatomical template (see (Makropoulos et al. 2018) for a description of the structural pipeline), as well as a resting-state functional MRI (rsfMRI) scan. A specialized 32-channel phased-array head coil was used to acquire the data (Hughes et al. 2017). To obtain rsfMRI scans, echo-planar imaging, with a multiband acceleration factor of 9 was used, with a TR of 392 ms and a TE of 38 ms (flip angle 34 degrees). Volumes had an isotropic resolution of 2.15 mm and we acquired 2300 volumes, for a total duration of 15:03.5 minutes.

### 2.3 Preprocessing and Quality Control

Data was processed using the minimally preprocessed pipeline from the dHCP, which has been described elsewhere (Fitzgibbon et al. 2020). Briefly, the dHCP pipeline strives to add minimal preprocessing, while performing rigorous quality control, improving signal to noise, and minimizing artifacts. As an overview, the pipeline included four major steps. First, the susceptibility distortion field was estimated. Second, the rsfMRI images were registered to the structural images in native space and then with the dHCP-40 week template. The third included motion correction, estimation of motion regressors, and dynamic susceptibility distortion correction. The final stage was a denoising step to reduce artifacts from multiband acquisition, cardiorespiratory effects, motion, and arteries (sICA). This step included FMRIB’s ICA-based Xnoiseifier, which identified noise components and then regressed them from the data (Salimi-Khorshidi et al. 2014). Before data release, the dHCP performed quality control, as described in the release notes (developingconnectome.org/data-release/second-data-release/release-notes). This resulted in the 512 infant datasets included with the release and 464 included in this study.

### 2.4 Tissue Iron Calculation

Tissue iron is paramagnetic and therefore, has an impact on the T2* signal. Indices of iron were obtained by normalizing and time-averaging T2* weighted images obtained during rsfMRI, which quantifies relative T2* relaxation rates across the brain (Brown et al., 2014; Langkammer et al., 2010; Larsen and Luna, 2015; Peterson et al., 2019; Price et al., 2021; Schenck and Zimmerman, 2004). Differentiating our analysis from fMRI BOLD studies, we are interested in aspects of the T2*-weighted signal that do not vary across time and have been shown to reflect properties related to tissue iron (Larsen & Luna, 2015). We first normalize each volume in the time course to the whole brain mean. This normalized signal is then aggregated voxelwise across all volumes using the median, which results in one image per participant that estimates the normalized T2* (nT2*) signal. High motion TRs with FD>0.3 were censored and excluded from analyses.

### 2.5 Regions of Interest

Bilateral regions of interest (ROIs) were taken from the FSL atlases. These included the putamen, and include all voxels combined across both hemispheres caudate and pallidum. Both the caudate and the putamen were identified using the Oxford-GSK-Imanova Structural and Connectivity Striatal atlas (Tziortzi et al., 2011). The region of interest for the pallidum came from the Harvard-Oxford subcortical atlas (Desikan et al., 2006). Two overlapping voxels were identified, coded as belonging to both the pallidum and putamen. These were assigned to the pallidum, which had fewer voxels.

In order to map the ROIs from adult MNI space into the dHCP-40 week template space, we first skull stripped the 40-week template, and used Advanced Normalization Tools (ANTs), shown to be effective in pediatric studies, to obtain a warp. Warping the adult template directly into the 0.5mm space resulted in substantial error, upon visual inspection. Thus, we resampled the dHCP-40 week template into 1mm space to obtain accurate mapping between images. To ensure that our ROIs were restricted to subcortical grey matter, we used the tissue type 7 (deep grey matter) to create a mask and disregarded voxels outside of the mask.

### 2.6 Linear Models

To model developmental change at the regional, univariate level, we extracted the median nT2* signal for each of the three regions of interest. Our use of a bilateral ROI enabled us to get a single value, comprised of the data from both hemispheres. To assess overall age-related relationships, we initially modeled nT2* with postmenstrual age (nT2*∼ pma). To test our primary hypothesis, we then modeled nT2* with gestational and postnatal age as separate terms in the model (nT2* ∼ gestational age + postnatal age). We ran this model separately for each of the three subregions. As premature birth has the potential to impact postnatal iron levels, we created a third model that categorized infants as preterm if they were born less than 35 weeks gestation and modeled the interaction between preterm birth and postnatal age (nT2* ∼ preterm + postnatal age + postnatal age*preterm). To see if this extended to a continuous interaction across the whole sample, we created a model with a continuous interaction term (nT2* ∼ gestational age + postnatal age + postnatal age*gestational age). These were repeated for each of the three basal ganglia subregions. To correct for multiple comparisons, we used a Bonferroni correction at the level of 6 tests (2 age variables x 3 ROIs). The significance cutoff for all analyses reported was p=0.008 (i.e. p<0.05/6). We report the coefficients from lm in R, modeled with the raw data.

### 2.7 Support Vector Regression

To model developmental change across the whole basal ganglia, multivariate level, we used epsilon support vector regression (SVR) with a linear kernel to predict age (postnatal or gestational) from voxelwise nT2* maps of the basal ganglia. To isolate the nT2* signal most strongly related to either gestational or postnatal age, we residualized at the voxel level for one age term, and used this to predict the other (e.g., we residualized the nT2* map for gestational age and predicted postnatal age from the residuals). Data was residualized for all N=464 participants in a single model.

Subregions may be differently predictive of age in term and preterm infants, so we created separate SVRs for infants less than 35 weeks (preterm) and greater than 35 weeks (term) gestation. SVR was implemented using libsvm in Matlab. SVR is commonly used in neuroscience, and, briefly, is less focused on minimizing error. Rather, it keeps the error within an acceptable range, instead focusing on finding an appropriate line (hyperplane) that fits the data to perform regression. Instead of minimizing the ordinary least squares, SVR focuses on minimizing *coefficients*, and the error is defined by a parameter, epsilon, that can be modified as an input to the model. To maximize the sample size, we trained our models with leave one out cross-validation and used the default values from the libsvm toolbox and a linear kernel.

## 3.0 Results

### 3.1 Linear Models: Postmenstrual Age (PMA)

To examine the overall association between age and nT2*, we first modeled tissue iron and postmenstrual age and report coefficients from lm in R, where we modeled the raw data. As described above, the significance cutoff for uncorrected p values was p=0.008. We report both the uncorrected and corrected (i.e. p*6) p values below. In the putamen, we found significant PMA-related increases (B=-0.0029, p_uncorrected_=0.00029, p_corrected_=0.0017). There were no significant relationships with PMA in the caudate (B=0.0013, p_uncorrected_=0.088, p_corrected_=0.528) or pallidum (B=-0.0006, p_uncorrected_=0.068, p_corrected_=0.408). These results are in **Figure 1**.

**Figure 1.**
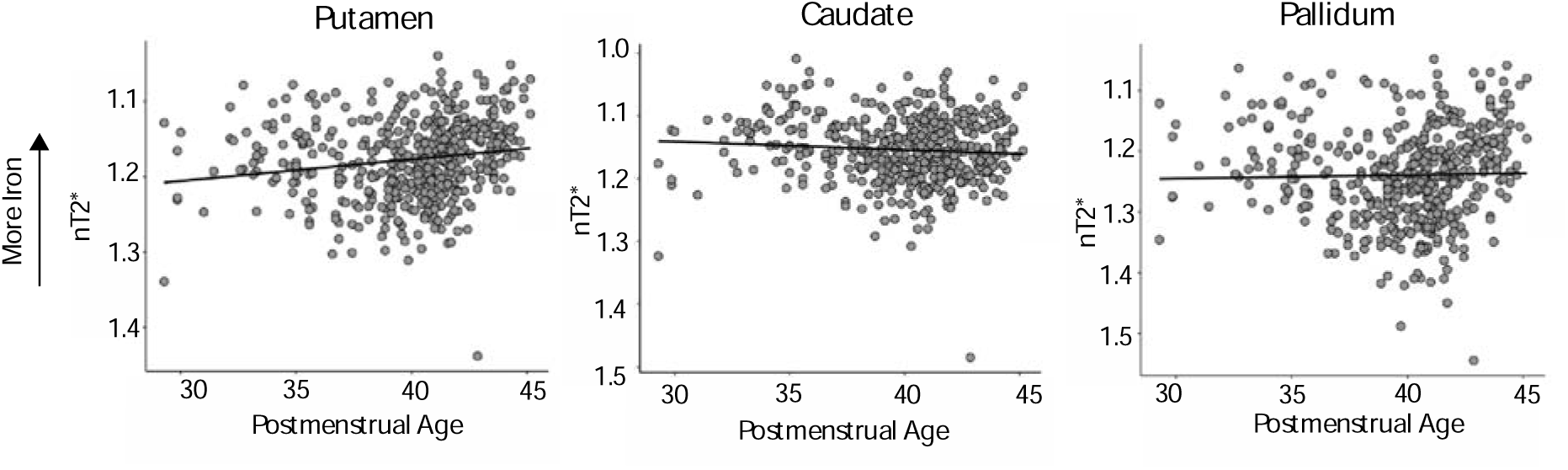
Postmenstrual age plotted with nT2* values in the putamen, caudate, and pallidum. Significant increases in postmenstrual age were associated with increases in nT2* in the putamen, but not the other basal ganglia subregions. nT2* is inversely related to iron content, thus Y-axes are inverted for visualization purposes.

### 3.2 Linear Models: Gestational and Postnatal Age

Gestational age and postnatal age were modeled as separate terms in linear models (nT2* ∼ gestational age + postnatal age). After correcting for multiple comparisons, gestational age was not associated with nT2* in the putamen (B=-0.0016, p_uncorrected_=0.02, p_corrected_=0.12), caudate (B=0.0019, p=0.013,) or pallidum (B=0.0017, p_uncorrected_=0.081, p_corrected_=0.486) (Figure 2A). Increased postnatal age was associated with increased nT2* in the putamen (B=-0.0109, p_uncorrected_=2e-16, p_corrected_=1.2e-15) and the pallidum (B=-0.0148, p_uncorrected_<2e-16, p_corrected_<1.2e-15), but the relationship between caudate nT2* and age did not survive multiple comparisons (B=-0.0023, p_uncorrected_=0.025, p_corrected_=0.15). These results are depicted in **Figure 2**.

**Figure 2.**
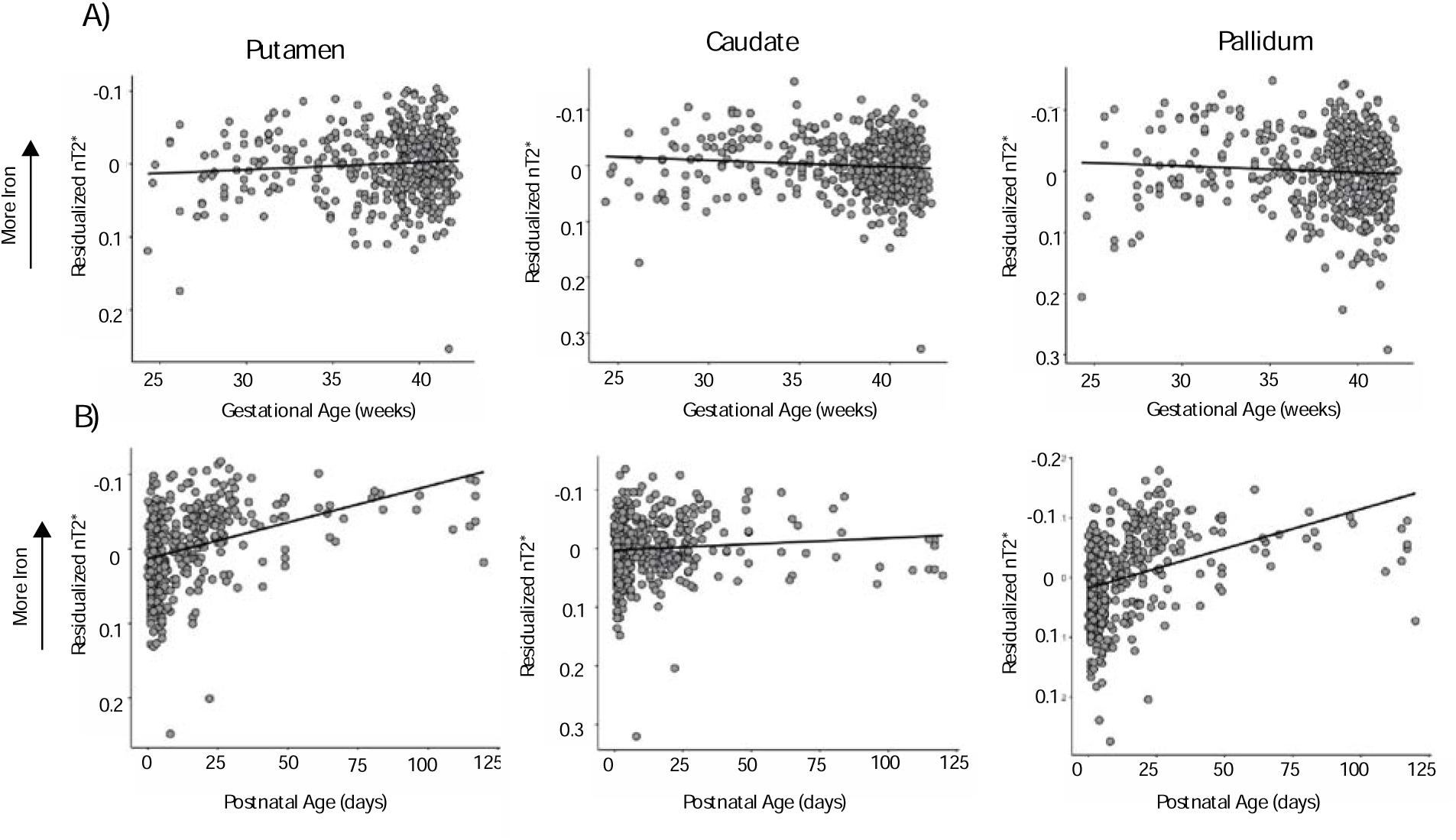
Age plotted with residualized nT2* for visualization. A) Gestational age plotted with nT2*, residualized for postnatal age. B) Postnatal age plotted with nT2*, residualized for gestational age. There were no significant relationships with gestational age in any of the three basal ganglia subregions. There were significant relationships with postnatal age in the putamen and pallidum, but not in the caudate.

### 3.3 Interactions Between Gestational Age and Postnatal Age

Decomposing early age variables allows us to accurately isolate the contribution of postnatal experience, which appears to be driving developmental effects. However, development in the postnatal period is likely partially dependent on prenatal variables, such as the iron stores that are acquired during pregnancy but may only be used in combination with experience in the postnatal period. To assess the impact of prematurity on postnatal development, we tested whether there was a categorical interaction between preterm birth, categorized as a gestational age of <35 weeks (n = 84), and postnatal age (nT2* ∼ preterm + postnatal age + postnatal age*preterm). In the putamen (B=0.0096, p_uncorrected_=2.72e-07, p_corrected_=1.632e-06), caudate (B=0.0063 p_uncorrected_=0.00213, p_corrected_=0.01278), and pallidum (B=0.0165, p_uncorrected_=5.19e-11, p_corrected_=3.114e-10) the interaction terms were significant in the model, demonstrating the interaction between preterm birth postnatal age was related to iron deposition. The relationship is plotted in **Figure 3** where, overall, premature infants started with higher iron levels at the time of scan than the term infants, but cross-sectional age-associations for iron were less evident in comparison to the term infants. To support this, we examined the main effect of prematurity in the interaction model described above. In the pallidum (B=-0.0594, p_uncorrected_=8.06e-09, p_corrected_=4.836e-08) and caudate (B=-0.0336, p_uncorrected_= 7.25e-05, p_corrected_= 0.000435), preterm infants had higher iron than term infants. In the pallidum (B=-0.020666, p_uncorrected_=0.0179, p_corrected_=0.1074) and caudate (B=-0.0188851, p_uncorrected_=0.00706, p_corrected_= 0.04236) there was a main effect of prematurity in a model without the interaction term, but this did not survive a correction for multiple comparisons in the pallidum.

**Figure 3.**
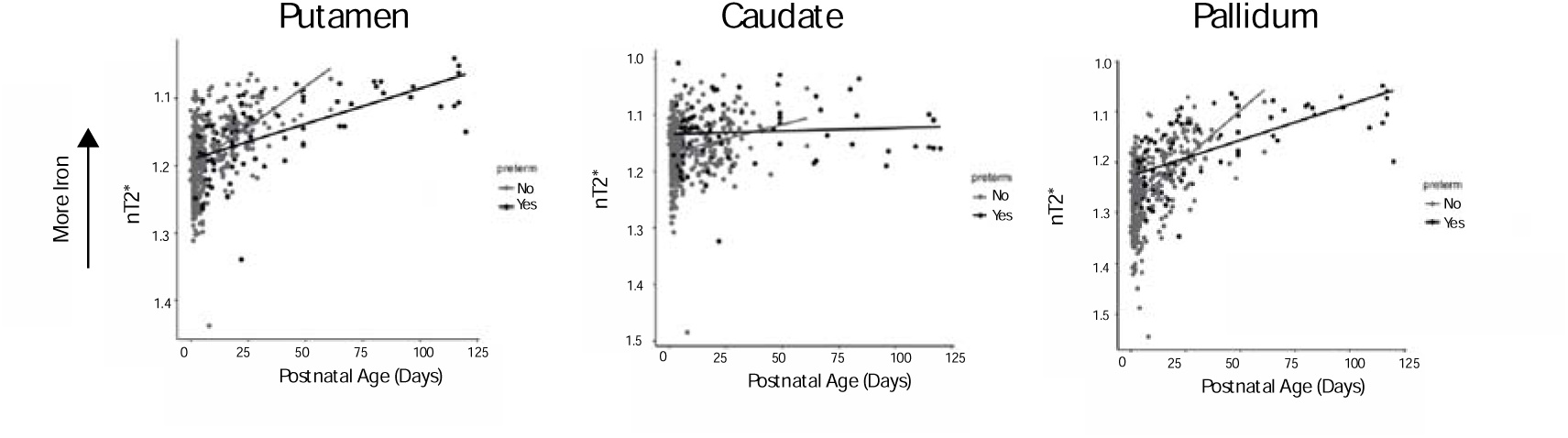
Postnatal age and nT2* plotted with term and preterm (<35 weeks gestation) infants in the putamen, caudate, and pallidum. nT2* is inversely related to iron thus the y axis is inverted for visualization. All three regions had significant interactions between prematurity and postnatal age.

It is important to assess if the term and preterm infants were matched for postnatal age. To address this, we did two things. First, we used the MatchIt package in R (version 4.5.4), with nearest neighbor interpolation, to identify the term infant that had the closest postnatal age to every preterm infant. To get the most accurate matches, we matched on both postnatal age and sex. As there are N=84 term infants, we were left with a sample of 168. There was still a significant interaction in the pallidum (B=0.008600, p_uncorrected_=0.0229, p_corrected_=0.1374), putamen (B= 0.0065007, p_uncorrected_ =0.0219, p_corrected_= 0.1314), and the caudate (B=0.007212, p_uncorrected_=0.02380, p_corrected_= 0.1428). The results do not hold up to a correction for multiple comparisons. These results are depicted in **Supplementary** Figure 1.

The preterm infants in the dHCP have greater postnatal ages than the term infants. To address this, while being mindful of reduced sample size, we removed 15 preterm infants with the greatest postnatal ages, keeping any infant with a postnatal age less than 65 days. This allowed for close term comparisons, with the largest term postnatal age being 61 days. Due to the reduced sample size (N=69 preterm), we did not match these infants (which would have resulted in a total sample of 138) and instead chose to consider the full 449 remaining neonates. There was still a significant interaction in the pallidum (B=0.0085, p_uncorrected_=0.01912, p_corrected_=0.11472) and putamen (B= 0.0060, p_uncorrected_=0.0274, p_corrected_= 0.1644), but not the caudate (B= 0.0039, p_uncorrected_=0.19587, p_corrected_>1). The interactions in the pallidum and putamen did not hold up to a correction for multiple comparisons.

Assessing the categorical interaction between term and preterm infants does not assess the continuity of the effect within the full sample. To do this, we created a model with a continuous interaction term using gestational and postnatal age (nT2* ∼ gestational age + postnatal age + gestational age*postnatal age). The pattern of significance was the same, with infants with shorter gestational ages starting off with higher iron levels and changing less over time. The interaction was significant in the pallidum (B=-0.0015, p_uncorrected_=7.81e-14, p_corrected_=4.686e-13), the putamen (B=-0.0008, p_uncorrected_=1.17e-07, p_corrected_=7.02e-07), and the caudate (B=-0.0005, p_uncorrected_=0.003789, p_corrected_= 0.022734).

### 3.4 Quadratic Models

In an exploratory analysis, we assessed if quadratic models fit the data better than linear models, assessed with the AIC() function in R. To achieve this, we mean centered the postmenstrual and gestational age terms in all models, but we did not do this for postnatal age, as some infants were scanned the same day they were born, and there are meaningful values of 0.

Figure 1 depicts nT2* and postmenstrual age. In the putamen, the model with the lowest AIC was the quadratic model (AIC=-1435.477). However, this does not disentangle the contribution of gestational and postnatal age, depicted in Figure 2. We then modeled the data with quadratic terms for both postnatal and gestational age, finding the model that fit the data best was the model with quadratic terms for both gestational and postnatal age (AIC=-1577.677). However, this model had a similar AIC to the model with a quadratic postnatal age term and a linear gestational age term (AIC=-1577.677). When splitting the data into preterm vs term infants, the models that fit the data best were models with linear terms for gestational age and quadratic terms for postnatal age in both the term (AIC=-1285.729) and preterm groups (AIC=-285.5301). Scatter plots for the best models in the putamen are in **Supplementary** Figure 2, and a regression table for the models can be found in **Supplementary Table 1**.

In the caudate, we considered the same models. When examining postmenstrual age, we found that the linear model had the lowest AIC (AIC=-1435.028). When further adding quadratic terms for both gestational and postnatal age, we found that the model that best estimated nT2* was the model with quadratic terms for both gestational and postnatal age (AIC=-1470.203). Once broken into preterm and term groups, the model that best estimated nT2* for the preterm infants was the model with scan age as a quadratic term (AIC=-270.9127). For the term infants, a linear model has a better AIC (AIC=-1210.957). A regression table for the models is found in **Supplementary Table 2** and figures depicting the results are found in **Supplementary** Figure 3.

In the pallidum, we repeated the same analysis with postmenstrual age and found that the model with a quadratic term for postmenstrual age outperformed the linear model (Figure 1; AIC: – 1090.248). When adding models with quadratic terms for gestational and postnatal age to our model comparisons, we find similar results in the full sample (AIC=1314.135), term (AIC=-1062.1450) and preterm groups (AIC=-251.8652), with the model with a linear term for gestational age and a quadratic term for postnatal age outperforming as the best model. These results are in **Supplementary Table 3** and **Supplementary** Figure 4.

### 3.5 Overall SVR for gestational age

nT2* values for each voxel in the basal ganglia (putamen + caudate + pallidum) were residualized for postnatal age, leaving nT2* values that primarily reflected gestational age in all N=464 subjects. Models were then trained on term and preterm data (<35 weeks GA) separately. Predicted age was correlated with actual age to get a measure of accuracy. Here, we have changed our cutoff for statistical significance. As we have two age variables but only a single basal ganglia tissue iron map, we are correcting for p=0.05/2, and the threshold for significance is thus p=0.025. Corrected p values indicate p*2. GA could not be accurately predicted from either the term (r(378)=-0.02, p_uncorrected_=0.64, p_corrected_>1) or the preterm (r(82)=0.005, p_uncorrected_=0.96, p_corrected_>1) data (Figure 4A).

**Figure 4.**
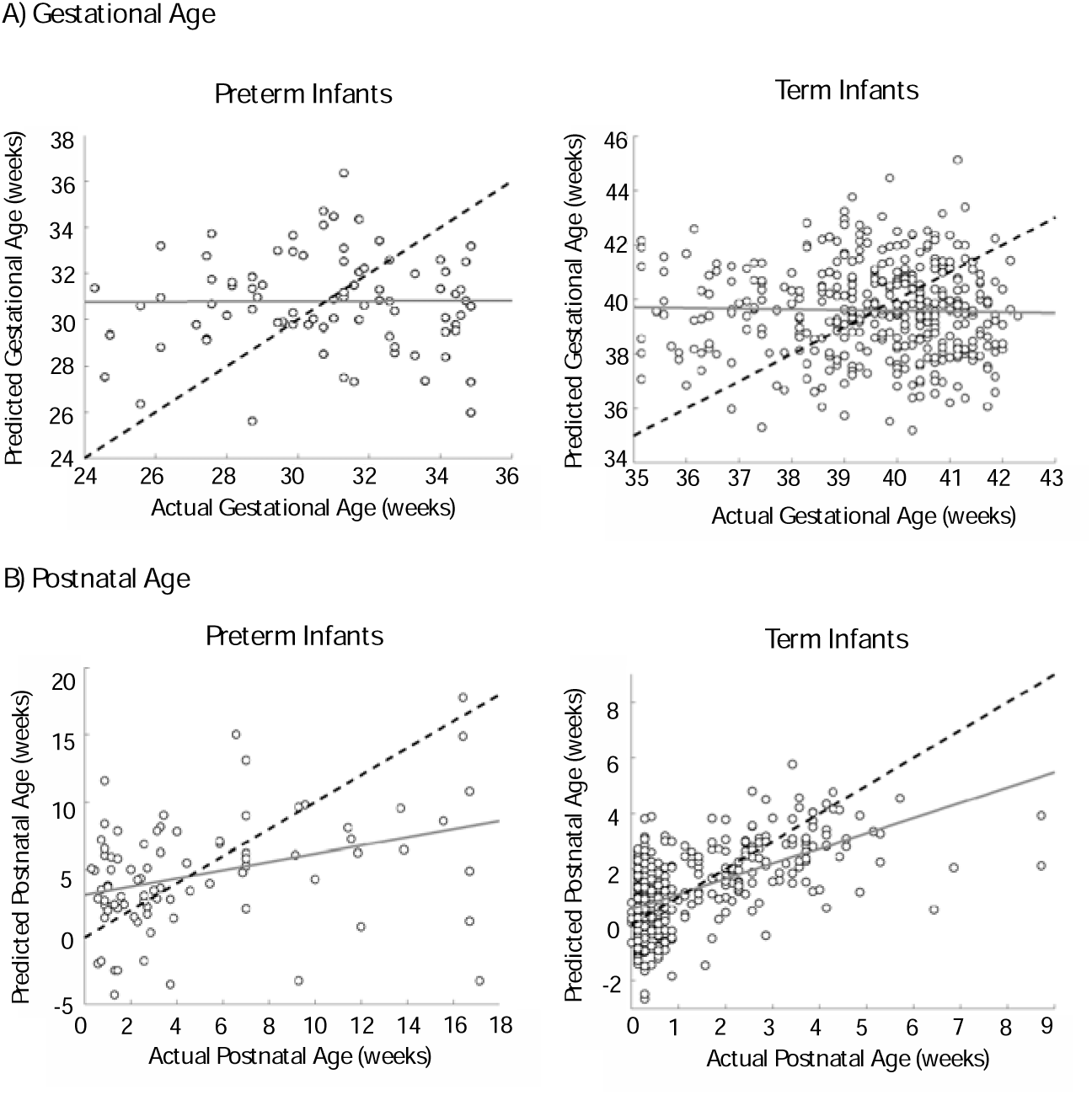
Scatter plots of predicted and actual age for the SVR models. A) Predicted and actual age values for models trained to predict gestational age from nT2* maps, residualized for postnatal age B) Predicted and actual age values for models trained to predict postnatal age from nT2* maps, residualized for gestational age. Black dotted lines depict what 100% accuracy between actual and predicted values would be, and grey solid lines are lines of best fit. Models were successful in predicting postnatal but not gestational age.

### 3.6 Overall SVR for postnatal age

nT2* values for each voxel in the overall basal ganglia ROI were residualized for gestational age, leaving values more strongly associated with postnatal age. Again, models were trained and tested on the term and preterm data separately. In contrast to the gestational age models, predicted age was correlated with actual age. Postnatal age could be predicted from both the preterm (r(82)=0.35, p_uncorrected_=0.0011, p_corrected_=0.0021) and term (r(378)=0.57, p_uncorrected_=1.0933e-34, p_corrected_=2.1866e-34) data (Figure 4B).

The preterm sample, N=84, was not large enough for k-fold cross validation, but the term sample N=380, was large enough to implement this. We used 5-fold cross validation with 304 neonates in each training group and an independent 76 for testing. Across the 5-folds, actual postnatal age was significantly correlated with predicted postnatal age (r=0.55, p_uncorrected_=2.4595e-07, p_corrected_=4.9190e-07; r=0.63 p_uncorrected_=1.2354e-09, p_corrected_=2.4707e-09; r=0.62, p_uncorrected_=1.8909e-09, p_corrected_=3.7818e-09; r=0.59, p_uncorrected_=1.5771e-08, p_corrected_=3.1541e-08; r=0.58, p_uncorrected_=3.9023e-08, p_corrected_=7.8046e-08). An exemplar plot from one of the 5 folds is in Figure 5.

**Figure 5.**
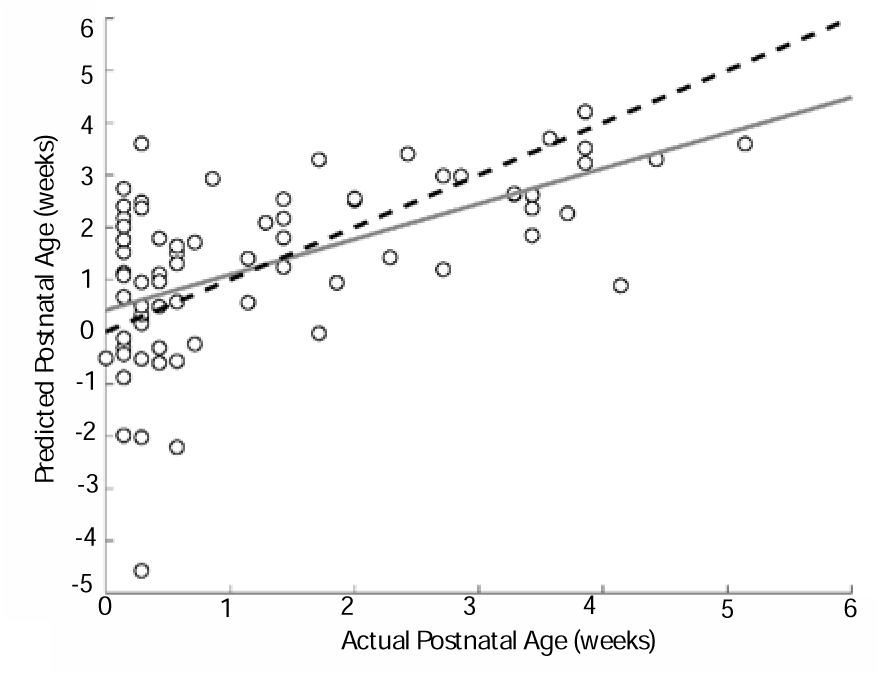
Scatter plots of predicted and actual postnatal age for an exemplar fold in the 5-fold cross validation analysis, where nT2* maps were residualized for gestational age and restricted to term infants. Black dotted lines depict what 100% accuracy between actual and predicted values would be, and solid grey lines are lines of best fit.

Although residualizing for postnatal age should remove data related to postnatal age from our voxelwise maps, there is the possibility that our models cannot account for all postnatal variability, and postnatal age could still be predicted from the residualized maps. To check if residualizing was removing a significant portion of the data related to postnatal age, we assessed if postnatal age could still be predicted from voxel-wise iron maps that had been residualized for postnatal age. Using leave one out cross validation, we found that even after residualizing for postnatal age, postnatal age could still be predicted from nT2* maps in both the term (r(378)=0.25, p_uncorrected_=1.2338e-06, p_corrected_=2.4675e-06) and preterm infants (r(82)=0.55, p_uncorrected_=1.2338e-06, p_corrected_=2.4675e-06)). This could be due to regional differences in iron deposition, where, even after postnatal age is accounted for on a voxelwise basis, regional differences between voxels could still allow for the prediction of postnatal age (i.e. iron deposition clusters in similar places with age). This possibility is returned to in the discussion.

However, predicting postnatal age from maps residualized for postnatal age still leaves in information about gestational age. It is also important to note that in this sample, gestational age and postnatal age are correlated. This is especially true in the preterm infants (r(82)= –0.65, p_uncorrected_=2.956e-11, p_corrected_=5.912e-11), but can also be seen in the term sample (r(378)= – 0.16, p_uncorrected_=0.001456, p_corrected_=0.002912). Preterm infants are often scanned later when they have lower gestational ages, as it may be difficult to scan them when they are younger and less stable. To explore this possibility that the correlation between postnatal age and gestational age was driving these effects, we extracted a portion of the term infant sample where gestational age and postnatal age were not correlated (all infants with a GA>37 weeks; N=343) and tried to predict postnatal age from nT2* maps where postnatal age had been residualized. Predicted postnatal age was still correlated with actual postnatal age, even after postnatal age had been residualized from the data (r(341)= 0.42, p_uncorrected_=2.8990e-16, p_corrected_=5.7979e-16), suggesting it is not a correlation between gestational age and postnatal age driving these effects. These results are depicted in **Supplementary** Figure 5.

#### 3.6.1 Spatial Distribution of Feature Weights

Using a linear kernel for SVR allows us to extract meaningful feature weights for each voxel from the trained model. SVR focuses on finding a function with the appropriate hyperplane to best fit the data to predict values of the dependent variable, y, from the model. An optimal SVR maximizes the distance between the hyperplane and data points, while minimizing error. The equation for the hyperplane becomes y=wx+b, where w is the weight, x is the data point and b is the bias. Thus, with a linear solution, the w for each feature becomes meaningful when compared with values of w from other features. It is important to note that this differs from linear regression, where the beta coefficients depict the strength of the relationship between the independent and dependent variable. Instead, values of w depict feature importance. Like in linear regression, values of w can be both positive and negative, and, can denote the direction of feature importance (Rodríguez-Pérez et al. 2017). In the case of our model, features with negative weights were important for predicting lower ages and features with higher weights for higher ages.

As a qualitative description of where the most highly weighted voxels were located (i.e. absolute weights, useful for examining a features overall importance in predicting age), we utilized the weights from the models that were separately trained on all of the term or preterm data (i.e. N=380 and N=84). We then identified voxels that had weights in the bottom or top 5% (10% highest overall), and then labeled them as belonging to the pallidum, putamen or caudate. This allowed us to see whether each region was highly weighted. As we are using one model per group, there are no significance tests between regions or groups. The distributions are in Figure 6. The distribution of weights is similar for both groups, with the caudate having the most voxels, expressed as a percentage of its total volume, while the putamen has the least. Despite the overall similarities between groups, there were still qualitative differences at the voxel level. An example spatial visualization for both groups is in Figure 7. A cluster of weights is seen on the head of the left caudate in both groups, while the medial portion of the caudate appears to be more highly weighted in preterm infants.

**Figure 6.**
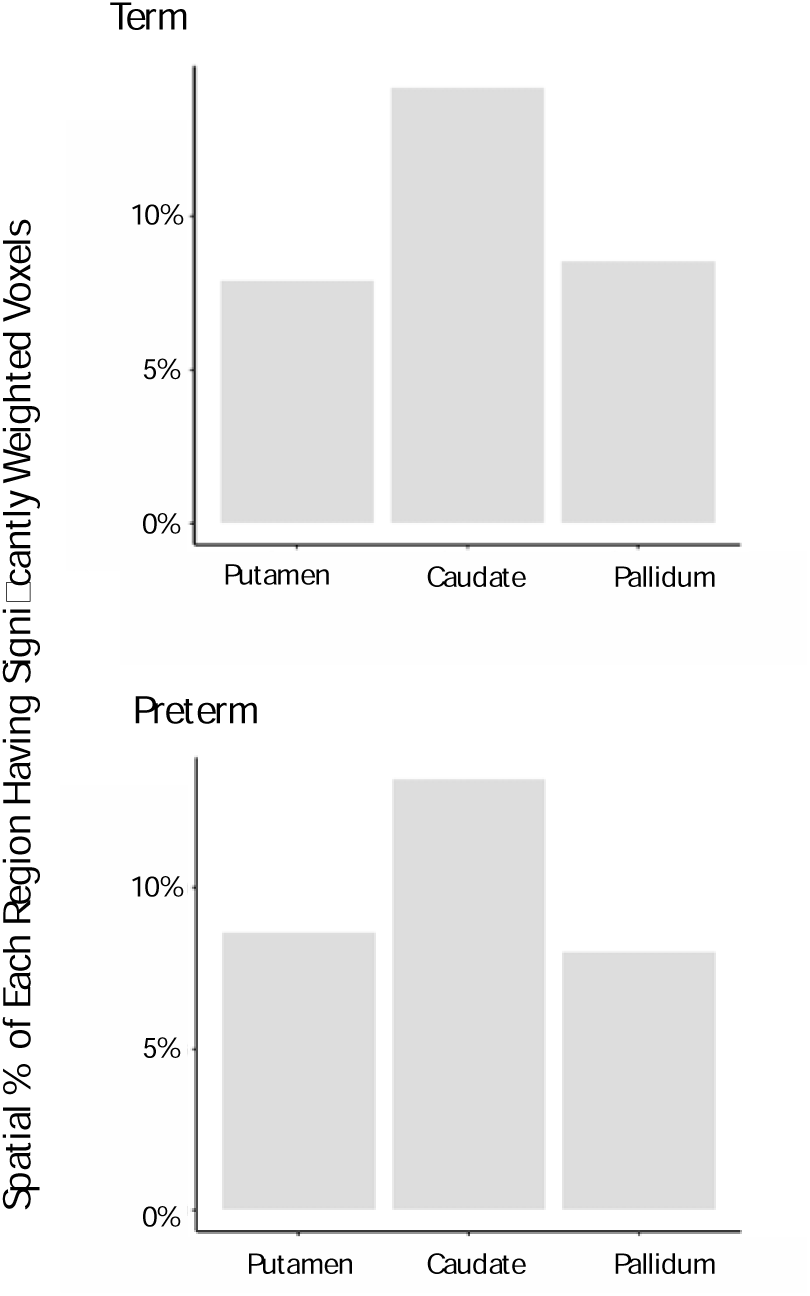
The spatial percentage of each region with feature weights that were the 10% most highly weighted. Bars represent the number of significantly weighted voxels, expressed as a percentage of region volume. The groups have weights that are roughly equally distributed.

**Figure 7.**
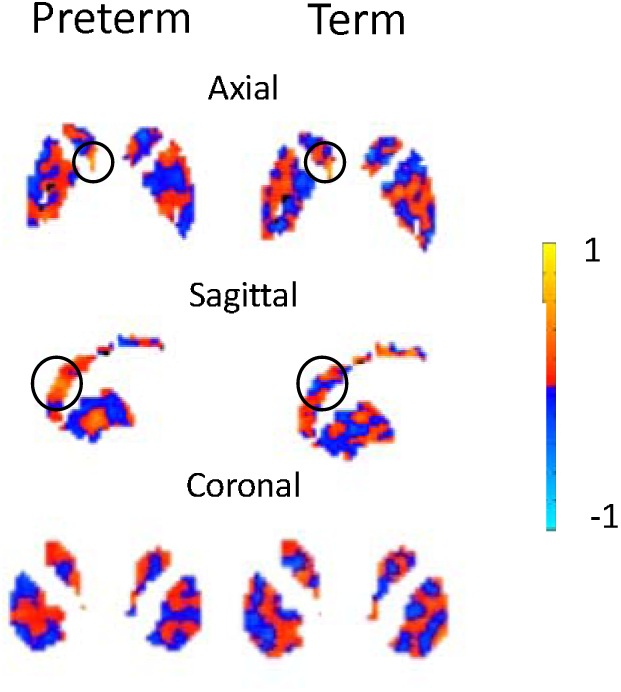
An exemplar of SVR feature weights rendered on the basal ganglia in A) preterm and B) term infants. Both groups have a small highly positively weighted cluster on the head of the left caudate (Box 1), but the preterm infants have more highly positively weighted voxels on medial portion of the caudate (Box 2).

To determine if the qualitatively clusters described above reached statistical significance, we used FSL’s threshold-free cluster enhancement (TFCE), going beyond regional t-tests, implementing it in the term infants because of the larger sample size. We chose TFCE specifically because this approach provides continuous sensitivity to the magnitude and spatial extent of effects in the data. Unlike traditional cluster-based methods that involve arbitrary cluster-forming thresholds, TFCE considers the entire range of data values, preserving information about subtle and widespread effects that might be missed by fixed thresholding methods. Traditional clustering methods often require the user to set an arbitrary cluster-forming threshold, which can be challenging to determine optimally. TFCE eliminates the need for such thresholding decisions, making the analysis less subjective and reducing the risk of false positives or false negatives, with the added benefit of improved statistical power. Finally, TFCE is robust to variations in noise levels and data smoothness. Traditional cluster-based methods may be sensitive to these factors, leading to variable results across different datasets or preprocessing pipelines. TFCE provides more consistent results across different data conditions. In order to generate independent samples of weights, we divided the N=380 into 19 groups of 20 infants, training a separate SVR for each. Weights were then extracted from the individual models, written back to image files, and fed into FSL’s TFCE as independent subjects in a one sample t-test. FSL gives each voxel a TFCE score, which indicates how likely it is to be a part of a local cluster. Values for each voxel were identified using a summary score based on the height and extent of the local signal. A cluster corresponding to the left caudate did reach statistical significance after a correction for multiple comparisons using family wise error correction. Voxelwise correction for multiple comparisons was implemented to deal with the significant number of voxels across the 19 models treated as subjects. Full results and visualizations are in **Supplementary** Figure 6.

## 4.0 Discussion

To determine significant, and potentially critical, periods of basal ganglia maturation in early development, we modeled the associations between basal ganglia tissue iron accrual and age, focusing on the unique contributions of gestational age, reflecting a neurobiological timetable and prenatal exposure, and postnatal age, reflecting additional influence from the environment. While we did not find associations with gestational age in any of the three basal ganglia subregions, we found strong associations with postnatal age, where increased age was associated with increased iron deposition. These results suggest that experience and sensory input post birth may be critical in the maturation of basal ganglia systems, reflected by the strong association with basal ganglia tissue iron.

Nevertheless, gestational age may not have been significant as a main effect in the model, but prenatal variables may still affect postnatal development. For example, the majority of an infant’s iron for the first 6 months of postnatal life is acquired in pregnancy (Rao and Georgieff 2007) (Dube et al. 2010), and preterm birth may decrease these stores by limiting access to the maternal environment (Cibulskis et al. 2021), influencing iron available for postnatal tissue iron accrual. Therefore, we tested if there was an interaction between preterm birth (or gestational age) and postnatal age. There were significant interactions in all three of the basal ganglia subregions with preterm infants starting off with higher iron levels in the caudate and pallidum but changing less over time than the term infants in all regions. In contrast with lower deposition that could be expected from lower iron stores, the increased deposition seen here might be reflective of the anemia common in premature infants, causing hypoxia, with the potential to lead to increased iron deposition and altered trajectories through oxidative stress (Kalteren et al. 2021).

Associations with age were stronger in some regions than others, and it is likely that examining age-related trends on a regional basis masks trends at a smaller spatial resolution. Clusters of voxels may mature more rapidly through their differential participation in feedback loops or connectivity with other regions. Iron localizes where myelin and dopaminergic processing (Greene et al. 2014) (Lao et al. 2016) are needed. The systems basal ganglia clusters are differentially connected with mature at different rates, with the potential to draw in iron to perform these functions at different developmental timepoints, which may be visible on a subregional level. These patterns may be disrupted based on preterm birth (Rose et al. 2008; Kanel et al. 2021). To address this, we used SVR to train models to predict postnatal age or gestational age from voxelwise maps of nT2*, residualized for the other age term, training separate SVRs for the term and preterm infants. We were able to significantly predict postnatal age in both the term and preterm groups, but, consistent with the main effects of the linear regression models, we could not predict gestational age in either group. Interestingly, we could predict postnatal age in the preterm infants despite the smaller sample size and larger range of ages.

Despite showing less change with age, the caudate had the most voxels, expressed as a % of volume, that were predictive of postnatal age in both the term and preterm groups, suggesting that subregions of the caudate change significantly with age, but this change might be masked using a regional approach. For example, both the term and preterm infants had a highly weighted cluster of voxels on the head of the left caudate, and additional clusters were not consistently visible on the body or tail. Clusters on the head of the caudate could be reflective of the subregions connectivity with orbitofrontal cortex (Fettes et al. 2017) and suggested participation within the striatal association loop (Gremel and Lovinger 2017). Frontal systems mature throughout development (Luna 2009), and the caudate’s role in these processes might be visible through subregional change with age. In contrast, the tail and body of the caudate may participate in the visual corticostriatal loop. Visual areas already show substantial maturity in infancy (Deen et al. 2017; Cabral et al. 2022), which could reflect the lack of change seen here. Taken together, when visualizing the overall distribution of weights, both groups had broadly similar patterns, potentially indicating that although the preterm trajectory could be slowed, as seen in the interaction models, it is not severely disrupted.

In the SVR analysis, we could still predict postnatal age after residualizing for postnatal age in our nT2* maps. The ability to predict postnatal age from data that has been residualized for it may be reflective of regional differences between iron deposition. Even after age is removed on a voxelwise basis, the differences between voxels may still be predictive of age. This is possible and would support the idea that iron accrues in clustered regions at different times through development, potentially indicative of functional specialization as motor, cognitive and reward systems come online. It is possible that the increased predictive ability when predicting postnatal age from data that has been residualized from it, rather than data that has been residualized for gestational age, is reflective of this relationship. Residualizing for postnatal age would leave in information about gestational age and voxelwise differences that may be predictive of postnatal age, both of which may be important in preterm development and result in better prediction accuracy.

Our results are in with those of Ning et al. (2019), who identified initial periods of linear change in older infants (3-6 postnatal months), finding linear relationships at 3 months in the pallidum and putamen, but not until 6 months in the caudate, supporting the protracted maturation we observed. Other groups have also measured postnatal change in older infants and toddlers, and found postnatal age effects in iron concentrations consistent with our results (Li et al. 2014; Ning et al. 2014; Zhang et al. 2018).

We showed that preterm infants have higher initial iron deposition in the pallidum and caudate. The initial increase in iron could be caused by iron deficiency and hypoxia (Peeples and Genaro-Mattos 2022). At birth, preterm infants experience high rates of anemia both because of decreased access to maternal iron stores (Cibulskis et al. 2021), but also because of blood loss associated with their clinical status or blood draws (Madsen et al. 2000). Anemia causes hypoxia, which causes the release of iron from stored tissue. If enough iron is released, it upsets the balance in the brain, causing iron mediated cell death (Viktorinova and Durfinova 2021). Importantly, iron mediated cell death results in lasting iron deposition, marking the damage to the tissue (Wu et al. 2019). The basal ganglia may be vulnerable to this process as it contains the majority of iron stored in the brain (Brass et al. 2006), but also because the products of iron mediated dopamine neurotoxicity have the potential to kill dopamine neurons during periods of critical change (Hare and Double 2016).

There are standard clinical guidelines that ensure preterm infants have iron supplementation after birth, ensuring severe anemia is transient (German and Juul 2021). However, in all three subregions, preterm infants cross sectionally changed less over time than the term infants did. It is possible that mild iron deficiency, or even differences between groups, causes less iron to be deposited. In early postnatal life, there is normative dip in hemoglobin, as red blood cells are produced, fetal red blood cells die, and the vascular volume expands (Rao and Georgieff 2007). In term infants, this process occurs between 6-8 weeks of postnatal age. In preterm infants, this typically occurs 1-5 weeks earlier (Cibulskis et al. 2021). It is possible that the preterm infants in the current study were experiencing this within the study period, but the majority of the term infants were not scanned within this timeframe. The observed difference in iron in the preterm infants could therefore result in less iron being deposited, deviating from a typical trajectory. Alternatively, slower overall iron accrual could occur as initial or transient hypoxic conditions may have killed enough neurons that iron-related storage and growth is no longer taking place with a normative trajectory.

The perinatal period represents a unique opportunity for postnatal stimulation, where new movements and feedback have the potential to influence basal ganglia development through experience and feedback from the environment. Motor, cognitive and reward systems work together to produce increasingly sophisticated behaviors. Infants as young as 10 weeks can be taught to kick to move a string attached to a mobile, which serves as a reward (Rovee and Rovee 1969; Rovee and Fagen 1976). As infants continue to respond to the environment, as described above, connectivity patterns and feedback loops have the potential to drive differential iron deposition by subregion. This has the potential to drive postnatal iron accrual in tandem with postnatal experience.

The pallidum interacts with other basal ganglia nuclei and sensorimotor areas through the motor portion of the basal ganglia-thalamo-cortical loop (Penney Jr. and Young 1986), and the initial maturation seen here could be because of initial motor development. However, the ventral pallidum is also thought to participate in reward learning and motivation (Tachibana and Hikosaka 2012). Early postnatal deposition could be driven by initial motor behavior, then as this behavior becomes reinforcing, the interaction between the two could begin the pallidum’s trajectory. Strong postnatal age effects were also found in the putamen, where motor development, through its participation in the motor cortico-striatal-thalamic-cortical loop, has the potential to drive iron deposition (Postuma and Dagher 2006; Smith et al. 2009). The putamen plays a role in initiating movements and in reversal learning, disengaging from ongoing actions, and these may be needed for the first time in postnatal life as infants interact with physical objects and actions influence bonding with caregivers (Groman et al. 2013; Haber 2016). However, in contrast, in our model that included a preterm by postnatal age interaction, the putamen was the only region that did not have a significant main effect of preterm birth, suggesting that overall iron deposition was not higher than term infants, indicating some preservation at the beginning of the putamen’s development.

We did not observe age-related effects in the caudate, after correcting for multiple comparisons. There was still a preterm by postnatal age interaction, but this did not remain significant after removing infants with a postnatal age greater than 65 days. The caudate is often implicated in cognitive functions, supported by studies in children looking at striatal iron deposition, which show iron accrual is related to working memory, spatial IQ, and general intelligence (Carpenter et al. 2000; Darki et al. 2016). The caudate is significantly connected to prefrontal regions, including the medial and lateral orbitofrontal cortex, and through this has been thought to participate in integrating sensory information, strategy or value-guided decisions, as well as other aspects of reward learning (Grahn et al. 2008; Fettes et al. 2017).

Additional roles for the caudate come from the participation of the tail in the visual corticostriatal loop, with significant connectivity to higher order temporal areas (Seger 2013). Iron deposition in the caudate could be influenced by early development of frontal regions, as supported by high weights, specifically in the head, in the SVR analysis. However, it could have a tempered trajectory driven by the maturation of visual areas. It is unclear if there another critical period of rapid growth as cognitive systems continue to come online.

Consistent with patterns seen here, the caudate has the lowest iron deposition in childhood (Hect et al. 2018) and adolescence (Larsen, Bourque, et al. 2020), relative to other striatal regions. A study of 1,543 youth ages 8-27 found the caudate to have significantly lower tissue iron than other basal ganglia ROIs (i.e., putamen, pallidum, and nucleus accumbens) (Larsen, Bourque, et al. 2020). Interestingly, the caudate exhibited the earliest stabilization in tissue iron (e.g., slowing of age-related change), mainly occurring at approximately 16. However, cognition was most closely associated with putamen iron in this sample. Elsewhere, in a slightly younger sample (ages 7 – 16), caudate iron was associated with both general intelligence as well as with processing speed (Hect et al. 2018). Taken together, these studies underscore the potential role of iron for cognitive development in later life (i.e., childhood and adolescence).

While we can speculate on the contribution of anemia and serum iron to basal ganglia tissue iron, future work should seek to link the two by acquiring serum iron and MRI in the same participants, at similar timepoints. As the majority of an infant’s iron stores are acquired prenatally (Cibulskis et al. 2021), maternal serum iron could also be quantified. Additionally, fetal MRI could characterize the relationship more fully. As iron is in breastmilk, and breastfed infants are already more likely to be iron deficient (Pizarro et al. 1991), a second blood draw could occur in the postpartum period. A limitation of our research is that the preterm infants had longer postnatal ages than the term infants. Although we attempted to account for this through our models, and, finally, by separating out preterm infants with the longest postnatal ages in analyses, future work should seek to balance this age variable among groups and extend the age of follow-up to investigate if preterm infants continue to change less. The relationship between neonatal tissue iron deposition and behavior should be characterized in infancy and through childhood. Large datasets, with consistent longitudinal follow-up, such as the Baby Connectome Project (Howell et al. 2019), would be useful to measure normative trajectories and link these to behavior. This should also be extended to clinical populations that may benefit, such as infants with congenital heart disease, who experience brain injury, hypoxia, anemia, and substantial behavioral deficits.

This work utilized a novel method to estimate brain tissue iron in neonates, indirectly quantifying nT2* by using rsfMRI. One limitation of using the T2* signal to quantify iron neonatally is that the T2* signal changes significantly in response to changes in brain water concentration. The amount of water in the infant brain decreases significantly in initial postnatal life, where brain water content decreases by 3% in the first 6 postnatal months (Smart et al. 1973; Blüml et al. 2013). Potentially compensating for this, calculating nT2* involves a normalization step, where voxels are normalized to an individual participant’s whole brain median nT2*. Therefore, each participant’s basal ganglia nT2* is relative to their developmental stage (and water content), allowing nT2* measurements to account for potential differences in water between participants. The nT2* values in the basal ganglia represent iron represent change above the whole brain nT2* in a volume. Future work should seek to extend the iron measurements acquired here by comparing them to a quantitative MRI-based iron measurement, like R2’. This would assess the similarities between iron values obtained from the different imaging sequences, which are significantly related in adolescence (e.g. R2’ and QSM) (Peterson et al.).

The present study results have implications for normative cognitive and motor development, demonstrating potential periods of critical change in distinct subregions of the basal ganglia. Change within critical periods, reinforced by experience, such as when an infant learns kicking will move a mobile through a string attached to their foot, may map onto later performance on cognitive and motor testing, and characterizing these relationships may help identify maladaptive trajectories. The basal ganglia is implicated across several psychiatric disorders involving abnormal cognition and reward processing, such as depression and bipolar disorder, and deviations from trajectories allow for potential early diagnosis and intervention. These findings may also be valuable for disorders that involve iron-related pathology like anemia and congenital heart disease. Importantly, the nT2*w iron estimates used here can be obtained in any dataset that uses T2*-weighted echo planar imaging, which is common in research fMRI protocols. This enables researchers to reconsider and reanalyze clinical datasets, with hard-to-reach populations that may be undergoing longitudinal follow-up. Extending the results here have the potential to contribute to the understanding of many unanswered questions in the development of basal ganglia physiology and broadly of motor, cognitive, and reward systems.

Taken together, these results provide new evidence suggesting that postnatal experience is critical to basal ganglia dopaminergic maturation. Postnatal trajectories, visible on a regional and subregional level, have the potential to be influenced by the duration of exposure to the prenatal environment. In sum, these provide support for further study of basal ganglia tissue iron in both perinatal and infant life, which will help to understand the development of motor, cognitive and reward systems.

