## Supplementary material for "Multivariate and Regional Age-Related Change in Basal Ganglia Iron in Neonates": Supp Figures

**Supplementary Figures**

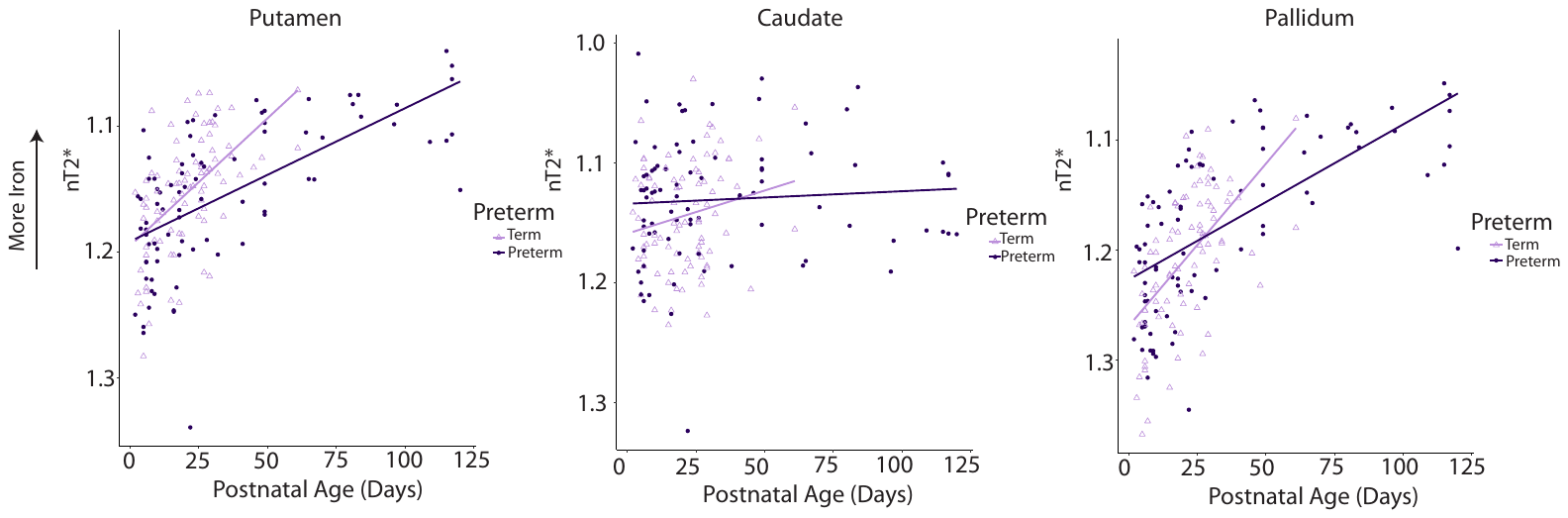

*Supplementary Figure 1.* Scatter plots for the three basal ganglia regions after using MatchIt to identify a matched term infant for every preterm infant. This was performed by matching for both sex and postnatal age. Term infants are scanned at younger ages than preterm infants in this dataset, and this helps to account for this. Significant interactions were found in all three subregions, but these did not hold up to a correction for multiple comparisons.

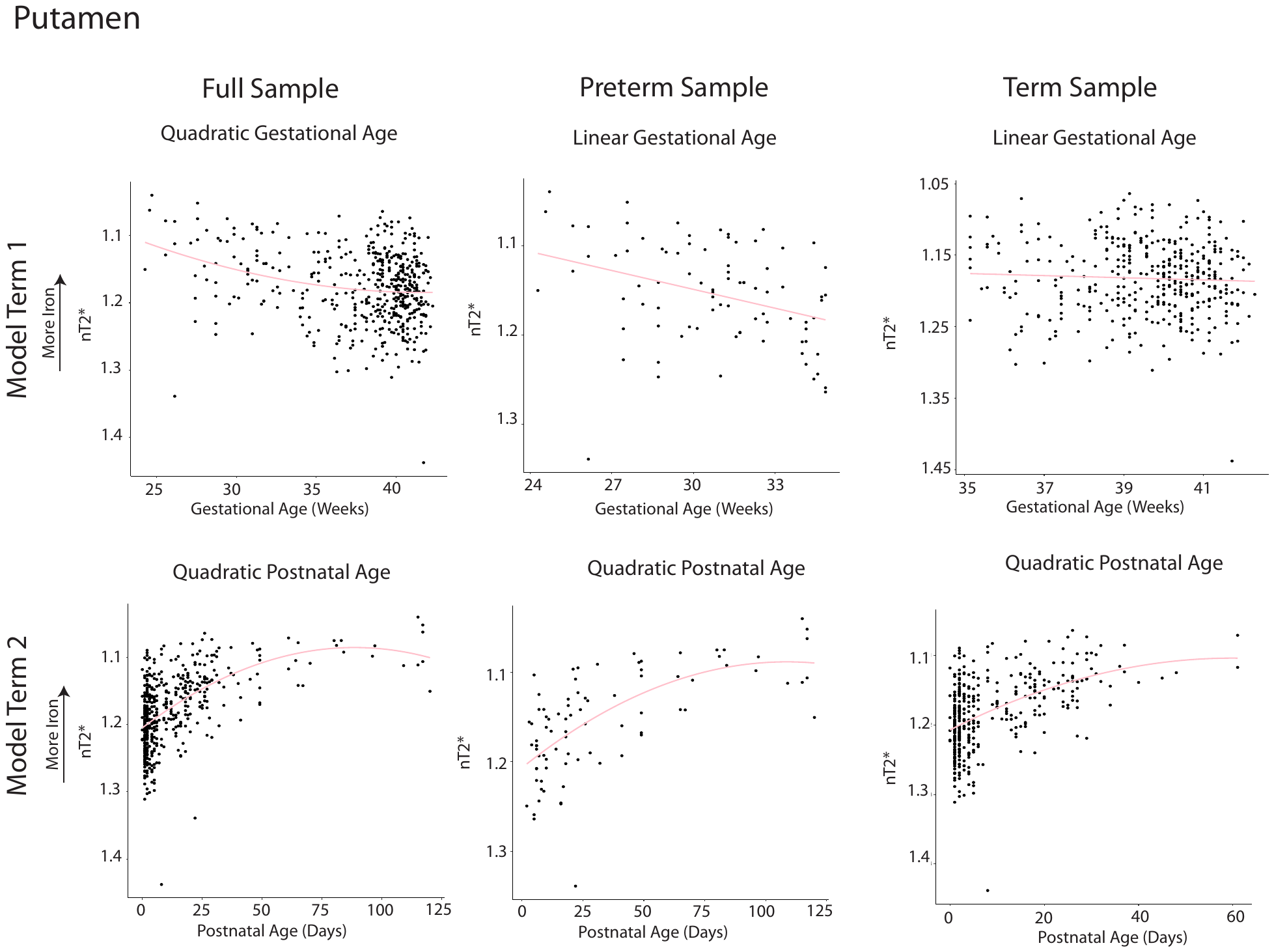

*Supplementary Figure 2.* Scatter plots depicting model terms for the full sample, preterm, and term infants in the putamen. In the full sample, a model with a quadratic term for gestational age (model term 1) and a quadratic term for postnatal age (model term 2) best fit the data. In the preterm infants, the model that best identified associations between nT2* and age was one with a linear term for gestational age (model term 1) and a quadratic term for postnatal age (model term 2). In the term sample, the model that best fit the data was one with a linear term for gestational age (model term 1) and a quadratic term for postnatal age (model term 2). Unlike in Figure 2, data plotted here is not residualized for gestational age or postnatal age before plotting, as this assumes a linear relationship.

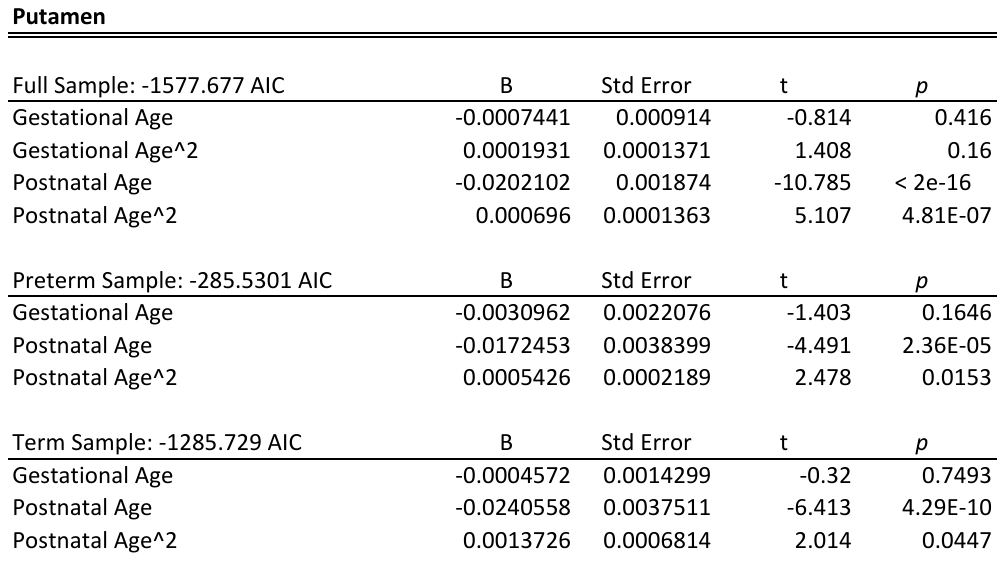

*Supplementary Table 1.* Regression table for the best model in the putamen (as selected by AIC) for each of the three groups. Coefficients were identified using lm() in R, where the raw data was modeled. Gestational age was mean centered in all models, but postnatal age was not as it contained meaningful values of 0.

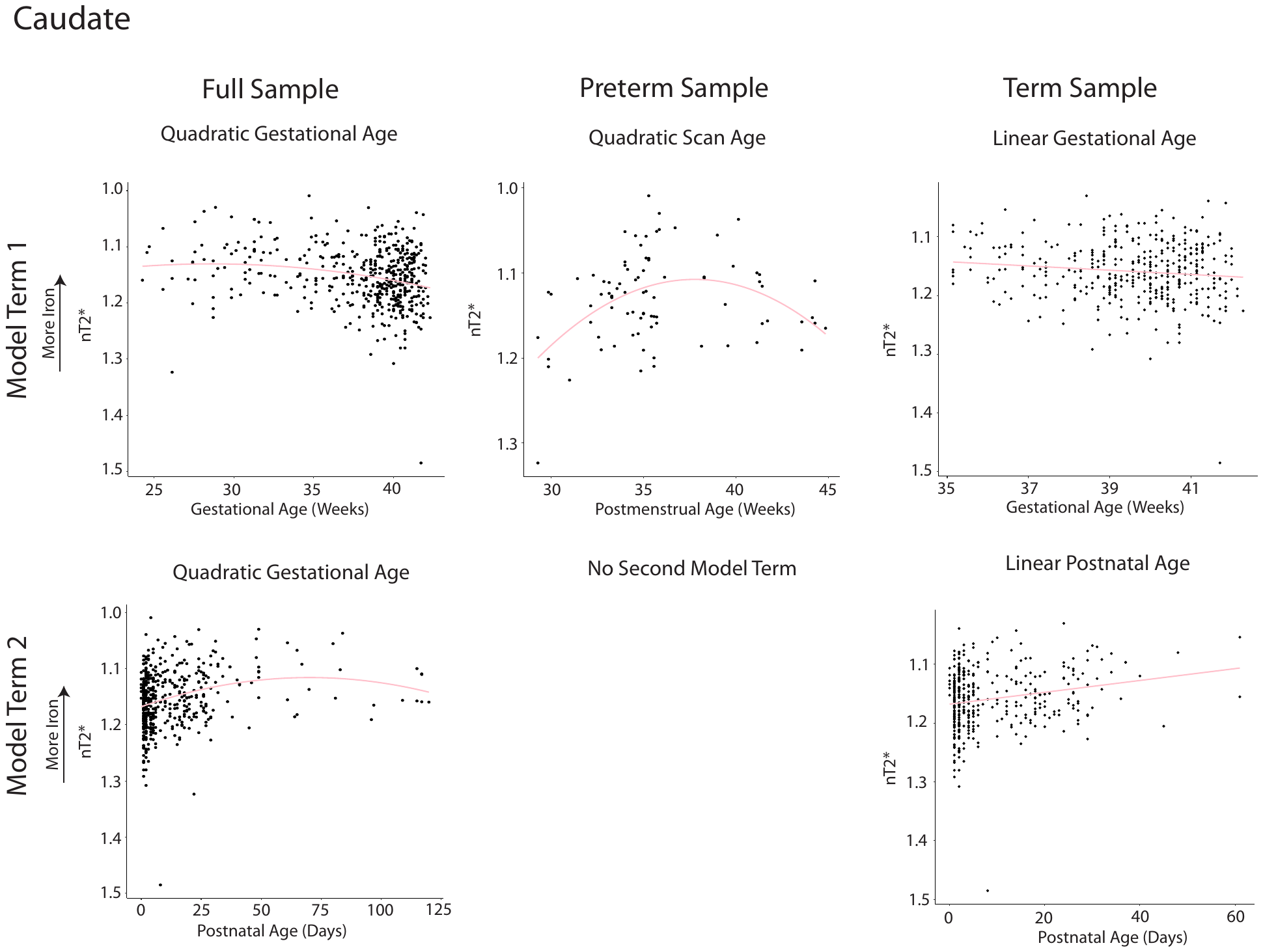

*Supplementary Figure 3.* Scatter plots depicting model terms for the full sample, preterm, and term infants in the caudate. In the full sample, a model with a quadratic term for gestational age (model term 1) and a quadratic term for postnatal age (model term 2) was the best fit. In the preterm infants, the model that resulted in the best association between age and nT2* was the model with a single quadratic term, scan age. In the term sample, this was best accomplished by a model with linear terms for gestational (model term 1) and postnatal age (model term 2), but this was not different from the model with two quadratic terms. Again, in contrast to Figure 2, data plotted here is not residualized for gestational age or postnatal age before plotting, as this assumes a linear relationship.

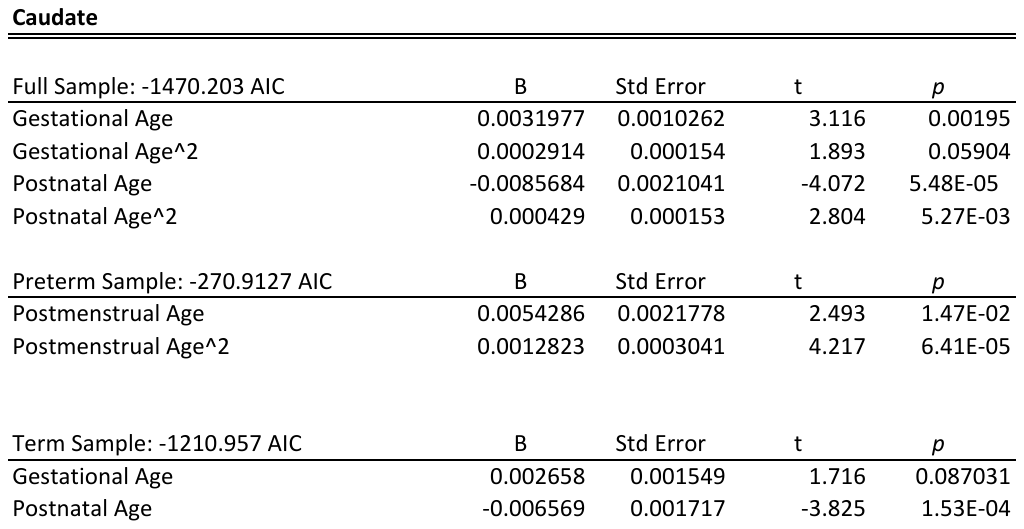

*Supplementary Table 2.* Regression table for the best model in the caudate (as selected by AIC) for each of the three groups. Coefficients were identified using lm() in R, where the raw data was modeled. Gestational age and postmenstrual age were mean centered in all models, but postnatal age was not as it contained meaningful values of 0.

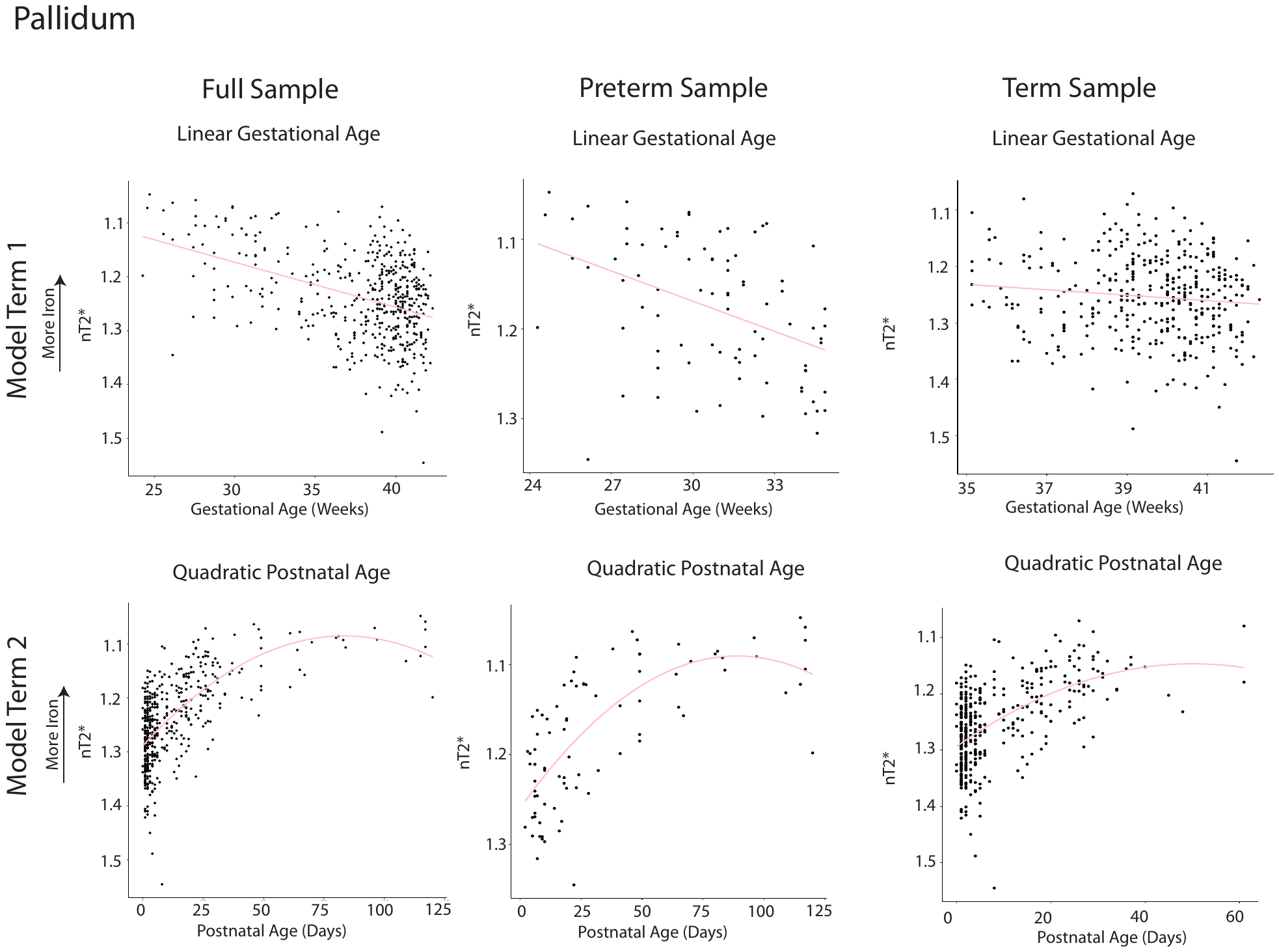

*Supplementary Figure 4.* Scatter plots depicting model terms for the full sample, preterm, and term infants in the pallidum. In the full sample, a model with a linear term for gestational age and a quadratic term for postnatal age was the best fit. This was also found for the term and preterm infants. Again, in contrast to Figure 2, data plotted here is not residualized for gestational age or postnatal age before plotting, as this assumes a linear relationship.

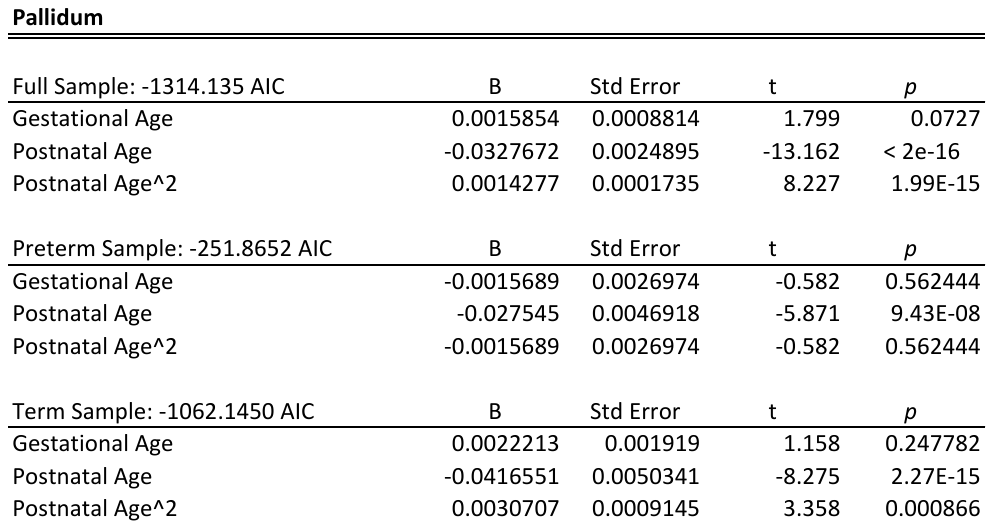

*Supplementary Table 3.* Regression table for the best model in the pallidum (as selected by AIC) for each of the three groups. Coefficients were identified using lm() in R, where the raw data was modeled. Gestational age was mean centered in all models, but postnatal age was not as it contained meaningful values of 0.

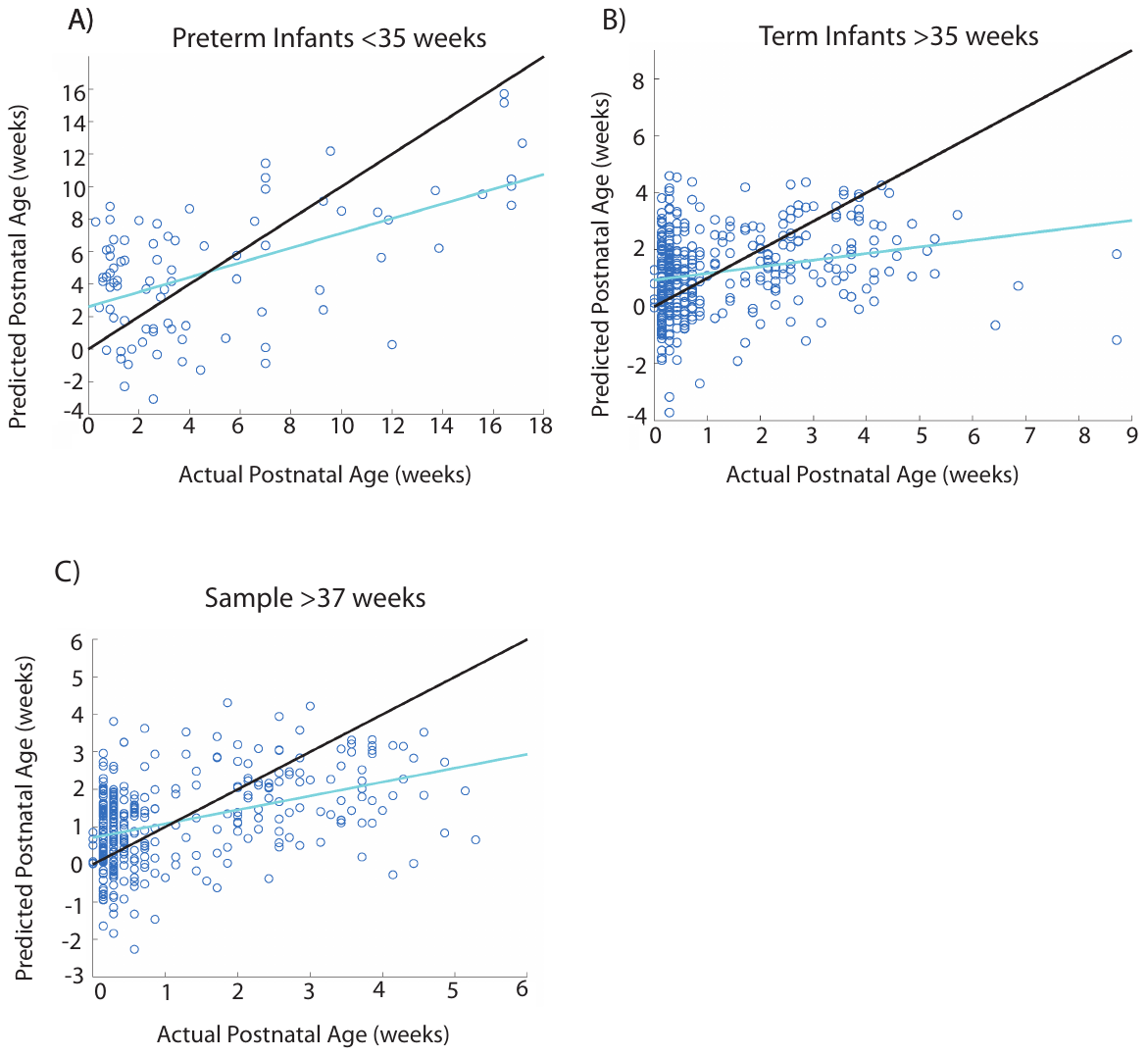

*Supplementary Figure 5* Scatter plots of predicted and actual postnatal age for the SVR models, after being residualized for postnatal age A) Predicted and actual age values for models for preterm infants <35 weeks gestation. B) Predicted and actual age values for models for term infants >35 weeks gestation. C) Predicted and actual age values for models with infants >37 weeks gestation where postnatal and gestational age are not correlated. Postnatal age could still be predicted from nT2* maps that had been residualized for postnatal age in all three groups.

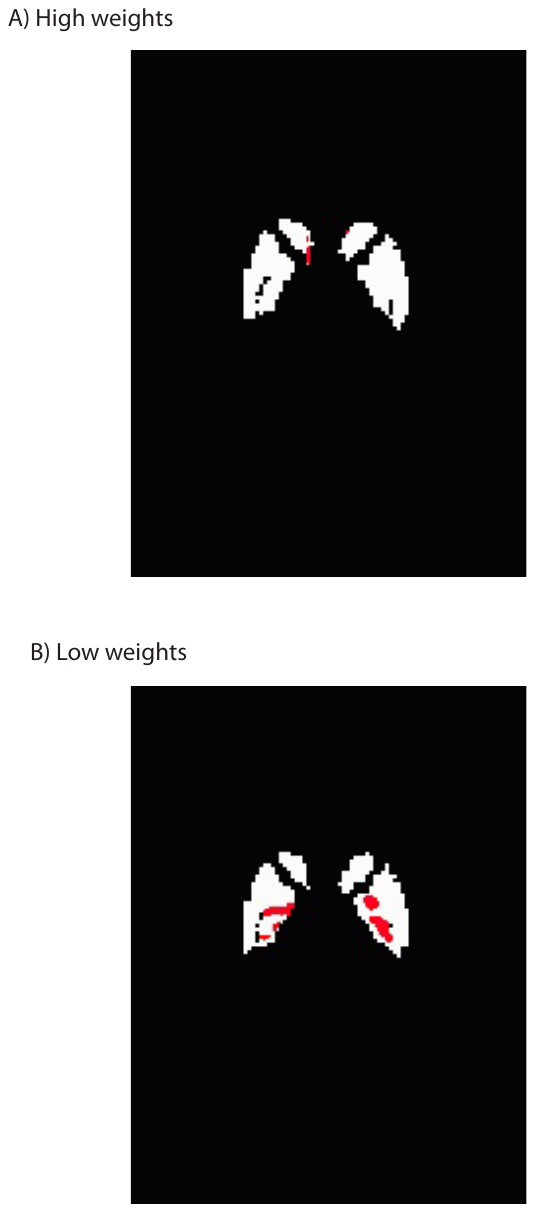

*Supplementary Figure 6.* Visualizations of the tfce clusters that were significant after FWE correction at p<0.05. For the positive weights (A) there was a significant cluster on the head of the left and right caudate, and for the negative weights (B) there were significant clusters that could be localized around the pallidum and putamen. To obtain p-values, the N=380 sample was split into 19 groups of 20, and separate models were trained for each group. This left 19 images, which we treated as “subjects” in a one sample t-test, run using FSL’s tfce’s to identify clusters.

The presence of significant clusters indicates regional nT2* associations occurring at distinct portions of development, implicating subsections of the three regions.
